# A standardized quantitative analysis strategy for stable isotope probing metagenomics

**DOI:** 10.1101/2022.12.20.521340

**Authors:** Dariia Vyshenska, Pranav Sampara, Kanwar Singh, Andy Tomatsu, W. Berkeley Kauffman, Erin E. Nuccio, Steven J. Blazewicz, Jennifer Pett-Ridge, Neha Varghese, Matthew Kellom, Alicia Clum, Robert Riley, Simon Roux, Emiley A. Eloe-Fadrosh, Ryan M. Ziels, Rex R. Malmstrom

## Abstract

Stable isotope probing (SIP) facilitates culture-independent identification of active microbial populations within complex ecosystems through isotopic enrichment of nucleic acids. Many SIP studies rely on 16S rRNA sequences to identify active taxa but connecting these sequences to specific bacterial genomes is often challenging. Here, we describe a standardized laboratory and analysis framework to quantify isotopic enrichment on a per-genome basis using shotgun metagenomics instead of 16S rRNA sequencing. To develop this framework, we explored various sample processing and analysis approaches using a designed microbiome where the identity of labeled genomes, and their level of isotopic enrichment, were experimentally controlled. With this ground truth dataset, we empirically assessed the accuracy of different analytic models for identifying active taxa, and examined how sequencing depth impacts the detection of isotopically labeled genomes. We also demonstrate that using synthetic DNA internal standards to measure absolute genome abundances in SIP density fractions improves estimates of isotopic enrichment. In addition, our study illustrates the utility of internal standards to reveal anomalies in sample handling that could negatively impact SIP metagenomic analyses if left undetected. Finally, we present *SIPmg*, an R package to facilitate the estimation of absolute abundances and perform statistical analyses for identifying labeled genomes within SIP metagenomic data. This experimentally validated analysis framework strengthens the foundation of DNA-SIP metagenomics as a tool for accurately measuring the *in situ* activity of environmental microbial populations and assessing their genomic potential.

**Importance:** Answering the question of ‘*who is eating what?’* within complex microbial communities is paramount for our ability to model, predict, and modulate microbiomes for improved human and planetary health. This question is often pursued using stable isotope probing to track the incorporation of labeled compounds into cellular DNA during microbial growth. However, with traditional stable isotope methods, it is challenging to establish links between an active microorganism’s taxonomic identity and genome composition, while providing quantitative estimates of the microorganism’s isotope incorporation rate. Here, we report an experimental and analytical workflow that lays the foundation for improved detection of metabolically active microorganisms and better quantitative estimates of genome-resolved isotope incorporation, which can be used to further refine ecosystem-scale models for carbon and nutrient fluxes within microbiomes.

## INTRODUCTION

The explosion of environmental sequencing data in the last decade has fueled a deeper understanding of the role of microbiomes in shaping human health, ecosystem function, and the Earth’s biogeochemical cycles (1). Further advancements in microbiome science require improved experimental approaches that link genomes to their *in situ* activities. Due to the limitations of culturing techniques, culture-independent methods that reveal *in situ* functions and link them to taxonomic identities play a crucial role in advancing the field of microbial ecology (2). Stable isotope probing (SIP) is a powerful cultivation-independent tool that links metabolic activity and taxonomic identity of environmental microbes (3). During a DNA-SIP experiment, compounds enriched with heavy stable isotopes (e.g., ^13^C, ^15^N, and ^18^O) are added to the microbial community of interest. The labeled compound is metabolized by active members of the microbial community and incorporated into cellular components, including DNA, during growth (4). As a result, the DNA of these active microbes becomes increasingly isotopically labeled, and, therefore, ‘heavier’ compared to the non-labeled DNA from inactive microbes (4). Isotopically-labeled DNA, referred to as ‘labeled’ from hereon, can be physically separated and recovered via isopycnic centrifugation using a CsCl gradient (5). Thus, microbes assimilating labeled compounds *in situ* can be identified through comparative sequence analysis of the DNA collected at different buoyant densities (BD) along the gradient.

Traditional DNA-SIP studies use 16S rRNA gene sequencing to identify labeled microorganisms (6, 7), and several analysis tools are available for 16S rRNA-based SIP studies (8-10). In addition to identifying microbial groups as either labeled or unlabeled, analysis tools such as quantitative SIP (qSIP) and delta BD (ΔBD) can also estimate the extent of isotope assimilation as atom fraction excess (AFE), which is the increase in the isotopic composition of DNA above background levels (11). Measurements of AFE can inform *in situ* growth rate estimates for specific microbial populations, enabling modeling of microbiome dynamics (12-14). Although 16S rRNA-based SIP analyses can taxonomically classify labeled microbes, the full genomic potential of metabolically active taxa are not always captured due to the difficulty in linking partial 16S rRNA gene sequences to their corresponding genomes (15). Adapting SIP analysis tools for the genomic level rather than the 16S rRNA gene level would enable genome-centric metagenomic SIP studies and establish stronger links between genomic information and *in situ* activity.

In recent years, multiple SIP studies have used metagenome sequencing in addition to, or in place of, 16S rRNA gene amplicon sequencing (16-21). We refer to this general approach as “SIP metagenomics” from here on to distinguish it from traditional 16S rRNA-based DNA-SIP. Some recent studies have applied the qSIP approach to shotgun sequencing data to estimate the isotopic enrichment of soil metagenome assembled genomes (MAGs) (22-24). While these represent exciting advancements in the field, SIP metagenomics faces challenges related to data analysis and interpretation. For example, estimates of isotopic enrichment depend on accurate measurements of absolute genome abundance, but determining genome abundance from metagenomic data is difficult due to its compositional nature (25-28). In addition, outstanding questions remain regarding optimal assembly strategies and the specificity and sensitivity of analysis tools given varying sequencing depth and genome coverage. Empirically answering these questions requires a defined experiment where the identity of labeled genomes and their level of isotopic enrichment is known *a priori*. To date, no such empirical study for validating SIP metagenomic sample processing and analysis has been published.

Here, we explore SIP metagenomic sample processing and analysis strategies using a designed microbiome where the identity of labeled genomes, and their level of enrichment, were experimentally controlled. We also investigated the utility of adding internal standards to monitor the quality of density gradient separations and normalize genome coverage levels. With this experimental design, we were able to: a) compare assembly methods for optimal genome recovery; b) determine how sequencing depth and genome coverage influence the detection of labeled genomes; c) examine how different approaches for measuring genome abundance impact estimates of AFE; and d) compare the sensitivity and specificity of different SIP analysis tools for accurately identifying labeled genomes. Based on our findings, we describe an experimentally validated strategy for SIP metagenomics and provide an R package (*SIPmg*) that adapts SIP analysis tools for shotgun metagenome sequence data, estimates absolute genome abundance within each fraction using internal standards, and identifies labeled genomes.

## RESULTS

To create a ground truth dataset for assessing SIP metagenomics, we generated a microbial community DNA sample where the identity of labeled genomes and their level of enrichment were known *a priori* (Fig. 1). Specifically, we combined unlabeled DNA extracted from a freshwater pond with aliquots of ^13^C-labeled *E. coli* DNA. We created eight levels of *E. coli* labeling ranging from 0 to 36 atom% ^13^C enrichment (Table S1). We also added two sets of synthetic DNA oligos at two different stages of sample processing to serve as internal standards (Fig. 1). The six “pre-centrifugation spike-in” standards had different BDs, each reaching maximum abundance in a different and predictable region of the density gradient (Table S2). Deviations from the expected distribution pattern indicated possible problems, such as a disturbance of the density gradient, that might compromise data quality from that sample (Fig. 2). The post-fractionation spike-ins, referred to as “sequins’’ hereafter (28) (Data Set S1), were added to each fraction after density separation (Fig. 1) to serve as internal calibration standards for calculating absolute genome abundances (Fig. 2). This experimental design provided a controlled dataset for answering questions regarding assembly strategies, genome abundance measurements, the impact of sequencing depth, and the accuracy of various SIP analysis methods.

**Figure 1.**
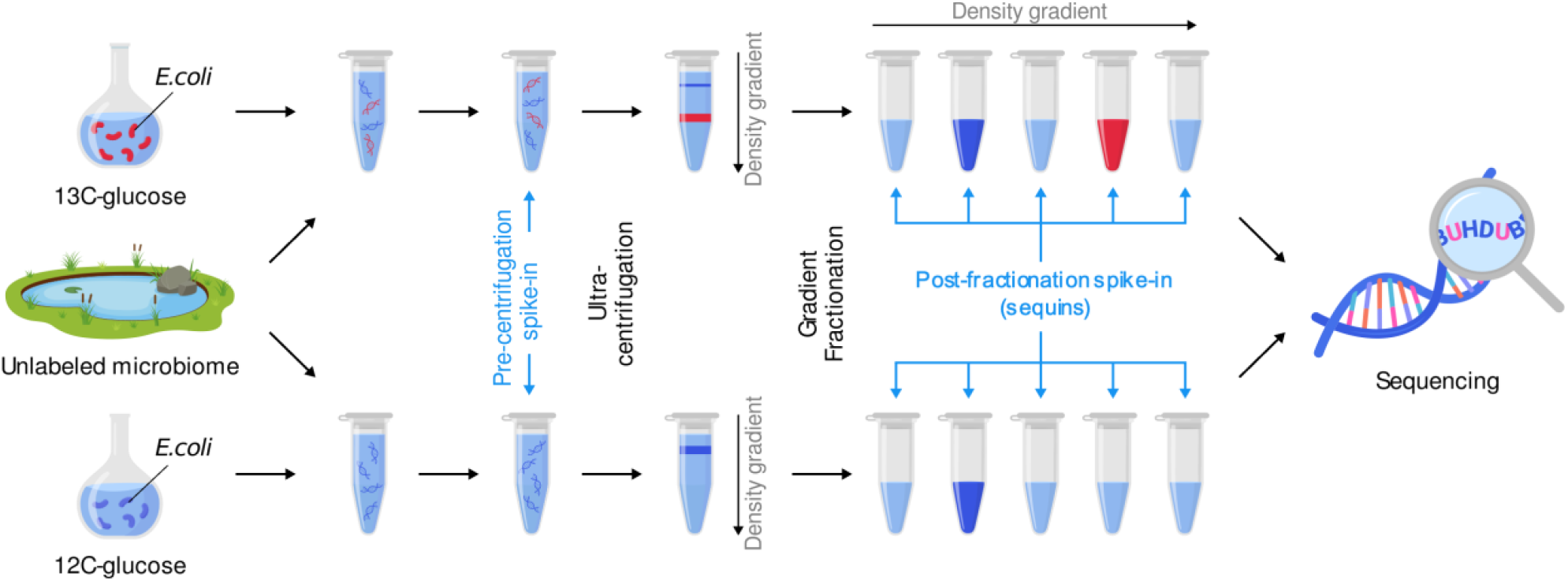
Experimental design and overview of laboratory steps in the SIP metagenomics workflow. To create a defined SIP experimental sample, DNA extracted from an unlabeled freshwater microbial community was amended with either labeled (^13^C) or unlabeled (^12^C) *E. coli* DNA. Pre-centrifugation spike-ins were added to each sample prior to ultracentrifugation in a CsCl gradient, and post-fractionation spike-ins (sequins) were added to each fraction after density gradient fractionation and collection. These two sets of synthetic DNA oligos served as internal standards to monitor the quality of density separations and normalize genome coverage levels.

**Figure 2.**
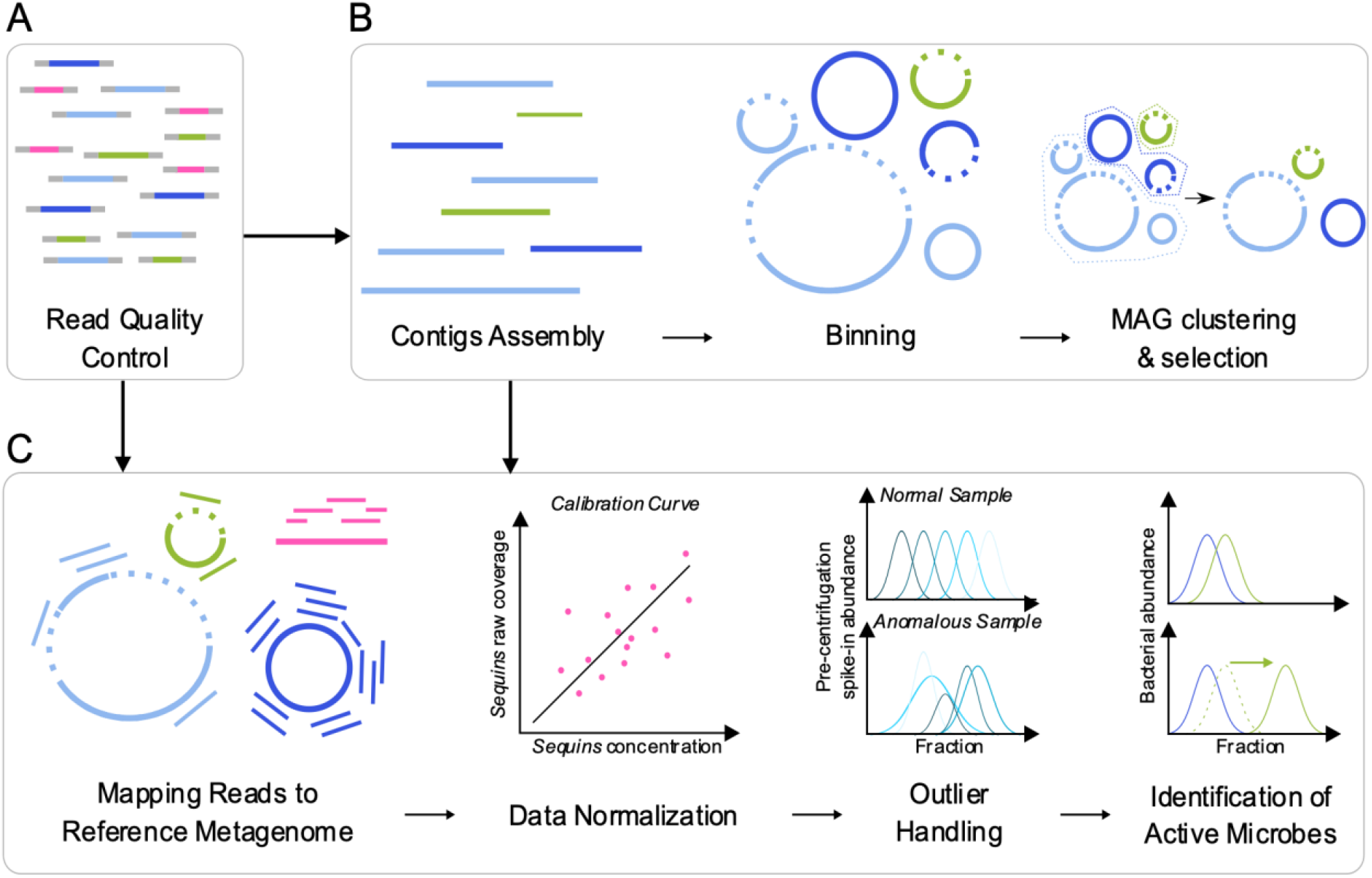
The workflow scheme for SIP metagenomic data analysis includes (A) quality filtering of the raw reads and (B) generation of a unique set of medium and high quality MAGs used for (C) quantification of absolute taxa abundances and identification of isotope incorporators. The addition of sequins provides the means for calculating absolute bacterial abundances (C, Data Normalization), and pre-centrifugation spike-ins aid in the detection of anomalous samples (C, Outlier Handling).

To develop an empirically validated workflow for SIP metagenomics, we next created the *SIPmg* R package, which was specifically designed to analyze shotgun sequence data from SIP studies. *SIPmg* calculates absolute taxon abundances using various methods, such as normalizing relative genome coverage to internal standards (this study) or total DNA concentrations (22, 23). *SIPmg* feeds taxon abundance into the HTS-SIP tool (29) where users can select different methods for identifying isotope incorporators, including qSIP (30), high-resolution SIP (HR-SIP, (8)), and moving-window high-resolution SIP (MW-HR-SIP, (9)). *SIPmg* also implements a version of the ΔBD method for estimating isotopic enrichment levels (8). To take advantage of metagenomic data, and similar to Greenlon et al. (23), *SIPmg* updates the qSIP model to use the observed GC content of assembled genomes rather than the estimated GC content used in qSIP analysis of 16S rRNA data (30). Finally, to correct for multiple comparisons, i.e. testing for significant isotope enrichment in multiple MAGs, *SIPmg* can adjust the confidence intervals around bootstrapped estimates of AFE using a variation of false discovery rate correction (31). With the *SIPmg* package, we evaluated the performance of different analysis approaches using our ground truth SIP metagenomics dataset.

### Maximizing recovery of metagenome-assembled genomes (MAGs) using individual and combined assemblies

In contrast to a typical metagenome sample, community DNA in a SIP experiment is separated into multiple fractions based on BD prior to sequencing (Fig. 1). Differences in GC content and levels of isotopic enrichment result in a non-random distribution of microbial genomes across the density gradient and sequencing each density fraction provides multiple options for assembly and binning. To determine the optimal strategy for maximizing MAG recovery, we compared assembly of the intact unfractionated sample, separate assemblies of each individual fraction, co-assembly of all fractions derived from the same initial sample, and a massive combined assembly using MetaHipMer (32) of all fractions from all samples. Each assembly was then independently binned using MetaBAT2 (33). A total of 2,022 MAGs were generated across all assemblies, of which 248 were high-quality, 447 were medium-quality, and 1,327 were low-quality as defined by the MIMAG reporting standards (34) (Data Set S2). The MetaHipMer assembly produced more MAGs than any other strategy. A total of 235 MAGs were recovered from the MetaHipMer assembly, of which 136 were medium- or high-quality (Fig. 3A). However, estimates of average MAG completeness and contamination for each assembly type were not substantially different (Fig. S1).

**Figure 3.**
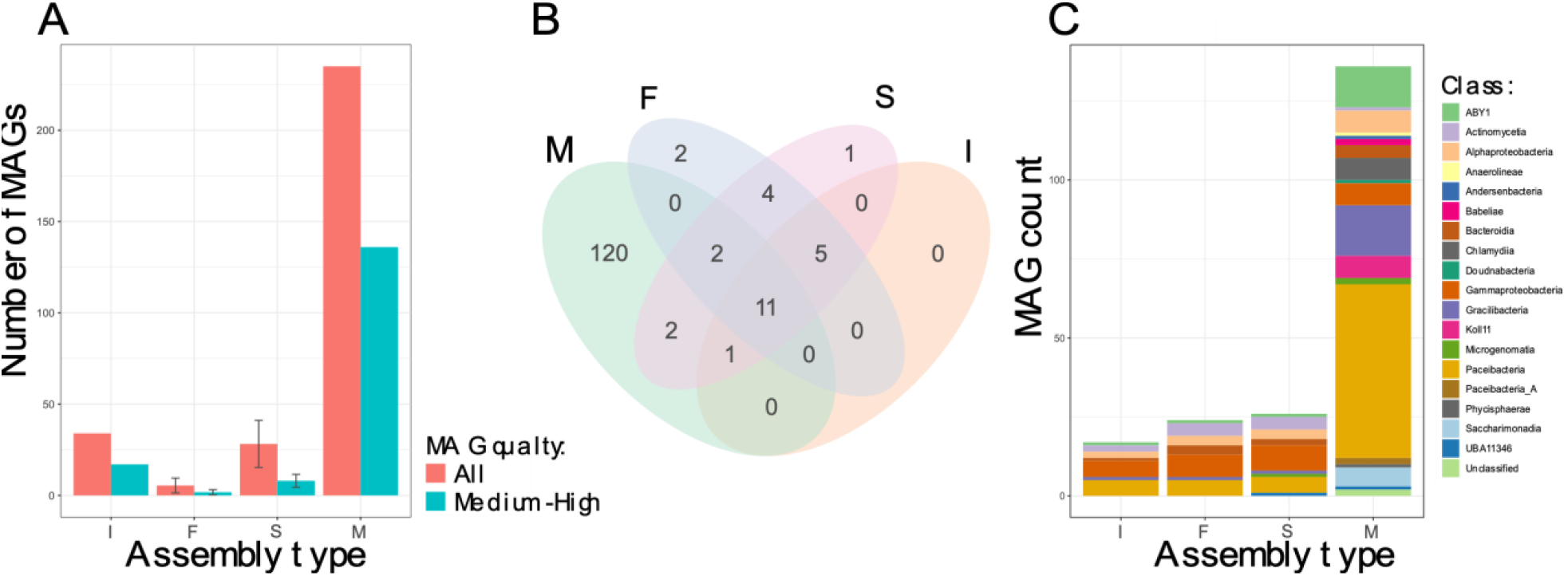
Comparison of metagenome assembly approaches for the SIP metagenome dataset generated from spiking *E. coli* into background unlabeled DNA from a freshwater microbiome. (A) Average number of medium- and high-quality MAGs recovered from different assembly approaches. (B) Venn diagram showing the number of unique and shared MAG clusters. (C) Compositional differences at the Class level recovered from different types of assemblies (I - intact metagenome assembly with MetaSPAdes, F - separate fractions assembled with metaSPAdes (n = 371 assemblies), S - all fractions within each replicate co-assembled with metaSPAdes (co-assembly of all fractions sequenced for a single SIP replicate sample, n = 24 assemblies), M - combined assembly of all fractions using MetaHipMer; for F and S the average number of MAGs was calculated, whiskers represent standard deviation across assembly type).

Next, we deduplicated all the medium- and high-quality MAGs recovered from all assemblies to determine whether any approach generated unique MAGs that were not present in other assembly types (Fig. 2B). We first grouped MAGs with average nucleotide identities of ≥ 96.5 and alignment fractions of ≥ 30% into a total of 148 unique clusters (35), then selected a single representative MAG for each cluster. Of these, 120 MAG clusters were exclusively produced by MetaHipMer. Twelve MAG clusters did not include any MetaHipMer-generated MAGs, and 11 of these clusters contained at least one MAG generated from the assemblies of individual fractions (Fig. 3B). Assembly of the intact unfractionated mock microbiome did not produce any unique MAGs (Fig. 3B). The different assembly strategies also produced MAGs with different taxonomic compositions. For example, MAGs derived from the MetaHipMer assembly accounted for an additional nine classes that were not present in other assemblies (e.g., *Anaerolineae, Andersenbacteria, Babeliae, Chlamydiia*, among others) (Fig. 3C). Most MAGs that were unique to the MetaHipMer co-assembly had lower coverage than MAGs recovered by other assembly approaches (Fig. S2). This suggests the MetaHipMer co-assembly captured more of the lower abundance MAGs in the samples than other assembly approaches, possibly due to the higher coverage levels that resulted from combining reads from all libraries (32). These results indicate that employing multiple assembly strategies and de-replicating the resulting MAGs can maximize genome recovery in SIP metagenomics studies.

### Anomalous sample detection using pre-centrifugation spike-in controls

As part of the quality control process, we devised an approach for detecting anomalous samples whose pre-centrifugation spike-in sequences displayed aberrant distributions along the BD gradient (Fig. 2C). We added six synthetic spike-ins to our samples prior to ultracentrifugation, and each spike-in had a different density based either on its GC content or the artificial introduction of ^13^C-labeled nucleotides during oligo synthesis (Table S2); therefore, each spike-in has a distinct and predictable peak in coverage along the BD gradient. Deviations from the expected spike-in distribution patterns may indicate events such as cross-contamination, library misidentification, or accidental disturbances of the density gradient significant enough to distort the distribution of MAGs throughout the gradient, all of which would introduce error into the downstream analysis. We identified three biological replicates with anomalous spike-in distribution patterns (Fig. S3), and these samples were removed from downstream analyses to avoid the introduction of extraneous noise. This example illustrates the utility of internal standards to illuminate quality control problems in SIP experiments that would otherwise go undetected.

### Normalizing genome coverage to quantify DNA isotope incorporation

Accurate abundance measurements are critical for determining levels of isotopic labeling. Briefly, models such as qSIP and ΔBD estimate a taxon’s AFE based on differences between its weighted BD in unlabeled controls and isotope-amended treatments (8, 30) (36), and weighted BD is calculated from the taxon’s abundance within each density fraction (see Methods equations 5 & 6). For amplicon-based qSIP studies, the relative abundance of a taxon is normalized to the total number 16S rRNA gene sequences within each fraction determined by qPCR (30). Estimating abundance in SIP metagenomic studies is more complicated, since shotgun sequencing lacks an equivalent method to 16S rRNA gene qPCR for absolute abundance scaling. Previous SIP metagenomic studies multiplied relative genome coverage with the total DNA concentration of each fraction (22, 23), which is a reasonable approach, although it does not account for potential variability introduced during DNA recovery, library creation, and sequencing of each fraction (27, 28, 37). By adding sequins to density fractions before DNA precipitation and recovery, we explored an alternative normalization strategy for measuring absolute abundance that could also account for variability in the downstream processing steps (22). In this approach, genome coverage within each fraction can be converted into absolute abundances through normalization based on the known concentration and observed coverage of the sequin internal standards. The AFE of each genome can then be estimated from these abundance measurements.

Our experimental design, where isotopic enrichment levels were known *a priori*, provided an opportunity to compare different approaches for calculating genome abundances and determine their impact on estimates of taxon AFE (Table 1, Fig. S4). More specifically, we compared the expected AFE values for labeled *E. coli* to AFE estimates from the qSIP model, with different approaches for calculating abundance, including: absolute abundance derived from normalization to sequins (Fig. 4A); absolute abundance estimated by multiplying either relative abundance or relative coverage with total DNA concentration (Fig. 4B and 4C, respectively); and relative coverage without conversion to absolute abundance (Fig. 4D). Results from all of the abundance normalization strategies we tested are provided in Fig. S4 and Table S3. Any genome other than *E. coli* that was identified as labeled was considered a false positive, whereas failure to identify *E. coli* as labeled was considered a false negative.

**Table 1:** Performance of different approaches for calculating genome abundance across density fractions based on the results from spiking ^13^C labeled *E. coli* DNA into background DNA of an unlabeled freshwater community. AFE was predicted using the qSIP model. Specificity was estimated as (true negatives)/(false positives + true negatives). Sensitivity was estimated as (true positives)/(true positives + false negatives)

| Method | Procedure | Specificity | Sensitivity | Spearman correlation between estimated & true AFE (*p*-values) |
| --- | --- | --- | --- | --- |
| Absolute abundance using sequins | Regression using sequin coverage and concentration | 0.993 | 0.857 | 0.85 (0.014) |
| Absolute abundance using total DNA concentration | Product of relative abundance and DNA concentration (23) | 0.991 | 0.714 | 0.8 (0.031) |
|  | Product of relative coverage and DNA concentration (22) | 0.922 | 0.571 | 0.27 (0.55) |
| Relative coverage | Relative coverage of MAGs in each fraction | 0.999 | 0.571 | 0.76 (0.046) |

**Figure 4:**
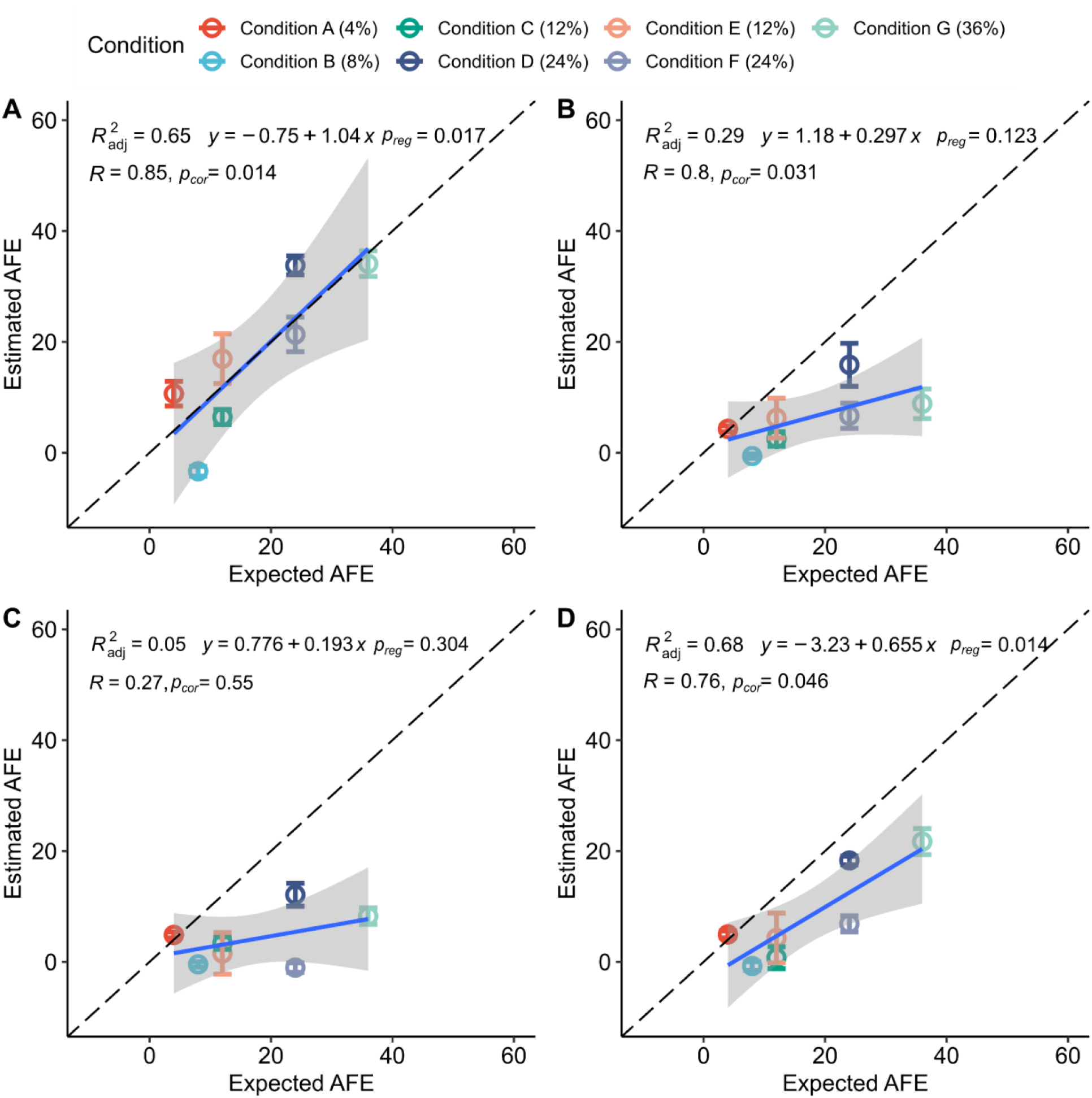
Comparison of predicted atom fraction excess (AFE) versus the expected AFE of *E. coli* using different approaches for measuring genome abundance across the density gradient. The qSIP method was used to estimate AFE in all cases. Genome abundance in each density fraction was determined by (A) normalization to sequin internal standards, (B) multiplying relative abundance with DNA concentration following Greenlon et al. (23), (C) multiplying relative coverage with DNA concentration following Starr et al. (22), and (D) relative coverage without additional normalization. For all comparisons, please refer to Table S3. Error bars represent the standard deviation of AFE calculated using the qSIP method’s bootstrapping approach. The expected AFE for each condition is in parentheses, and additional details about conditions, including replicate numbers, are provided in Table S1. *p*_*cor*_ and *p*_*reg*_ correspond to the *p-*values for the Spearman correlation and the linear regression F-statistic, respectively. The intercepts determined by linear regression were not significantly different from zero (*p-value* > 0.05) in any method for estimating abundance.

Abundance estimates derived from the sequin approach outperformed all other approaches based on combinatorial assessment of specificity (lower false positives), sensitivity (lower false negatives), and the Spearman correlation between expected and predicted AFE values (Fig. 4, Table 1, Table S3). The two approaches using total DNA concentrations did not produce statistically significant linear regressions (*p*-value > 0.05) between expected and estimated AFEs (Fig. 4B, 4C, Table S3), although the sensitivity for detecting labeled *E. coli* was the same or better than sensitivity using relative coverage (Table 1). Relative coverage produced the highest specificity, although it had lower sensitivity than the normalization approach using sequins (Fig. 4D and Table S3). These results suggest that internal quantification standards can improve estimates of genome abundance and AFE.

### Comparison of various SIP analysis method

In addition to qSIP, other analysis methods such as ΔBD (8), high-resolution SIP (HR-SIP, (8)), and moving-window high-resolution SIP (MW-HR-SIP, (9)) can identify labeled taxa. We compared all four approaches for their ability to accurately identify isotope incorporators in our defined SIP metagenomic dataset. We also compared estimates of *E. coli* AFE predicted with the ΔBD and qSIP methods; HR-SIP and MW-HR-SIP do not provide quantitative estimates of enrichment. For all methods, absolute genome abundances were determined by normalization to sequins.

The qSIP method predicted the level of AFE for *E. coli* with greater accuracy than the ΔBD method (Fig. 5). The qSIP approach also had higher specificity than the ΔBD method, producing only 7 false positives across all conditions compared to 12 false positives, respectively (Table S4). The MW-HR-SIP approach had the fewest false positives, with only 4 across all conditions, while maintaining the same sensitivity as the qSIP method (Table S4). The sensitivity and specificity of HR-SIP were lower than both MW-HR-SIP and qSIP methods (Table S4). Based on these results, we selected qSIP and MW-HR-SIP for further evaluation.

**Figure 5:**
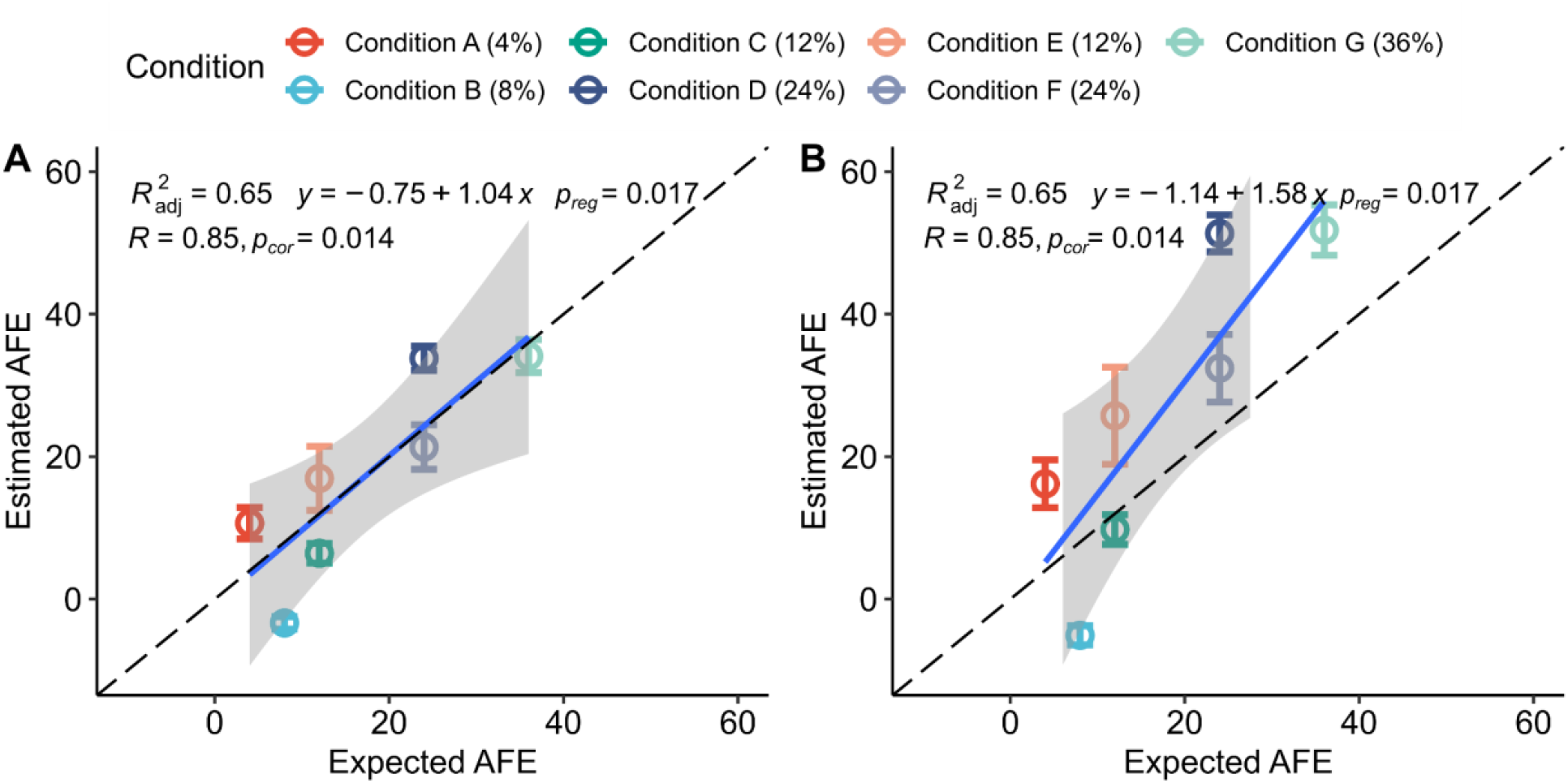
Comparison of AFE estimates produced by the (A) qSIP and (B) ΔBD methods using the mock metagenome where levels of *E. coli* isotopic enrichment were known *a priori*. Both of these methods used sequin-based normalization for estimating genome abundance. Error bars represent the standard deviation of AFE calculated using the qSIP method’s bootstrapping approach. The expected AFE of *E. coli* within each treatment condition is given in parentheses. *p*_*reg*_ and *p*_*cor*_ correspond to the *p-*values for the linear regression and Spearman correlation, respectively. The intercepts determined by linear regression for qSIP and AFE models were not significantly different from zero (*p-value* > 0.05).

### Lower limits of genome coverage for reliable detection of isotope labeling

Next, we examined how sequencing depth affected our ability to detect isotope incorporation. As demonstrated above, the accuracy of abundance measurements impacts the accuracy of AFE estimates, and these abundance measurements are derived from genome sequence coverage. The relative abundance of microbial taxa comprising complex communities can vary by orders of magnitude; thus, genome coverage within sequencing libraries can vary similarly (38). This suggests that AFE estimates might be less reliable for taxa with low coverage. To determine the lowest depth of coverage at which an AFE could be accurately estimated, we performed qSIP and MW-HR-SIP analyses after subsampling *E. coli* reads to 10%, 1%, 0.1%, 0.01%, and 0.001% of their initial levels (Table S5). In the respective subsampled datasets, *E. coli* had an average total coverage ranging from 0.01X to 1,400X coverage. Here, ‘total coverage’ refers to the cumulative coverage across all density fractions of an individual sample.

The qSIP model consistently identified *E. coli* as labeled when mean total coverage was ≥ 1X (Table S6). The correlation coefficient between actual and predicted AFEs was 0.8 within this coverage range (*p*-value <0.05; Fig. S6 and Table S7). However, at total coverages <1X, qSIP failed to detect *E. coli* as labeled in several experimental conditions, and the predicted AFEs were not significantly correlated to the expected AFEs (*p-*value > 0.05) (Fig. S6 and Table S7). The MW-HR-SIP method was also less sensitive at lower coverage levels, and at 100X mean total coverage, it only detected *E. coli* as labeled in 3 out of 7 experimental conditions (Table S6). These data suggest that estimates of isotope enrichment are less reliable in general when genome coverage is low.

### Strategies to improve accuracy of detecting isotopically labeled genomes

To improve the accuracy of SIP metagenomic analysis, we explored different strategies to reduce the number of genomes incorrectly identified as labeled (i.e., false positives). For example, the number of false negatives increased as coverage decreased; therefore we tested whether implementing minimum genome coverage requirements could reduce the number of false positives. Excluding genomes with mean total coverages <10X reduced the total number of MAGs analyzed from 147 to 113, and reduced false positives identified by qSIP from 7 to 4 without increasing false negatives (Tables S6 and S8). This improved the balanced accuracy from 0.925 to 0.927. Raising the minimum mean total coverage to 17X eliminated all false positives, yet reduced the number of remaining MAGs analyzed to 68. We did not test coverage limits for MW-HR-SIP because the method struggled to detect *E. coli* as labeled when coverage dropped below 100X (Table S6) and applying a threshold of 100X would have limited our analysis to only 17 genomes (Table S8). These results suggest that excluding genomes with low coverage can decrease false positives and increase balanced accuracy. Although the definition of “low coverage” will vary based on experimental conditions and individual assessments of the tradeoffs between sensitivity and specificity, these results also suggest that confidence in the identification of labeled genomes should decrease along with their coverage levels.

We also investigated if false positives could be reduced by implementing a minimum level of isotopic enrichment necessary for a genome to be considered labeled. That is, rather than simply requiring genomes to be significantly greater than 0% AFE, which is the default setting of the qSIP approach (30), we examined different minimum AFE thresholds ranging from 2% to 12.5% (Table S9). A genome was considered to be labeled if the lower bound of its AFE 95% CI was greater than the minimum AFE threshold. With AFE thresholds between 2% and 6%, total false positives dropped from 7 to 3 across all experimental treatments, but *E. coli* was no longer identified as labeled in one experimental condition. The balanced accuracy was also reduced from 0.925 without AFE thresholds to 0.856 with a 6% AFE threshold (Table S9). False positives were completely eliminated with a minimum AFE threshold of 12.5%, but sensitivity was so poor (0.286) that *E. coli* was only identified as labeled in 2 out of 7 conditions (Table S9). Minimum AFE limits could not be tested with MW-HR-SIP analysis because this method does not estimate levels of isotopic enrichment. Together, these results illustrate a trade-off between sensitivity and specificity when increasing the minimum AFE threshold above zero, and suggest that false positives can be reduced by increasing the AFE threshold at the potential cost of losing sensitivity for the detection of minimally labeled taxa.

The number and identity of false positives varied across SIP analysis methods, presumably due to differences in their underlying algorithms. Therefore, we hypothesized that the number of false positives might be reduced by taking the consensus of different analysis methods, i.e. requiring that two separate models predict a MAG is labeled. All false positive MAGs found in qSIP analysis were also false positives in ΔBD analysis, and thus taking the consensus of these two methods did not produce fewer false positives than qSIP alone (Table S10). In contrast, there was no overlap in the identity of false positive MAGs between the qSIP and MW-HR-SIP methods, and a union of their results completely eliminated false positives without producing any false negatives (Table S10). However, we found it more advantageous to apply MW-HR-SIP and qSIP sequentially rather than independently. MW-HR-SIP had greater specificity than qSIP, therefore it was used as a first-pass filter to detect putatively labeled genomes while minimizing false positives. This subset of putatively labeled genomes was then re-analyzed with the qSIP model. Only genomes first identified as labeled by MW-HR-SIP and later confirmed with a significantly positive AFE by qSIP were labeled. Applying the tools in series reduced the number of multiple hypotheses tested (e.g., MAGs tested for enrichment), which subsequently increased the statistical power for AFE estimation. That is, without the initial reduction in identified incorporators, the qSIP analysis would have otherwise included all MAGs in its statistical comparisons between treatment groups, resulting in a smaller *p-* value required for significance with multiple hypothesis testing. The increased statistical power obtained by applying the models in series resulted in tighter confidence intervals for the AFEs of *E. coli* (Table S11). These results indicate that using a combination of analysis tools can reduce false-positive detection, although the tools used and their order of application may vary depending on preferences for sensitivity versus specificity.

## DISCUSSION

DNA-SIP has been an established method in microbial ecology for many years and has primarily relied on 16S rRNA gene sequencing to identify active taxa (16, 30, 39, 40) (14). With decreases in sequencing costs and increases in compute capacity, DNA-SIP studies can now utilize shotgun metagenomic sequencing to establish links between population genomes and *in situ* activities (22-24, 41-43). In addition, automated sample preparation substantially increases the potential scale of SIP metagenomic studies and allows for more biological replication (24). However, the growth of SIP metagenomics also depends on adapting analysis tools to work with shotgun metagenomic data and validating their performance. To this end, we designed a mock SIP metagenome that enabled empirical testing of sample processing and data analysis strategies. Our results suggest some potential best-practices for SIP metagenomic studies that can serve as a foundation for future improvements.

Comparing assembly strategies for SIP metagenomic data was a key goal of our study. Previous SIP studies have used different strategies, including assembling unfractionated DNA, assembling individual SIP fractions, and co-assembling several fractions (22-24, 44, 45). However, it was not clear which assembly strategy produces the most medium- and high-quality MAGs. For instance, in computationally-simulated SIP experiments, the co-assembly of multiple fractions improved MAG recovery compared to the assembly of unfractionated DNA (45). In addition, the large amount of sequence data used in co-assemblies can recover rare genomes that would otherwise be lost due to insufficient coverage in smaller assemblies of individual datasets (32). Conversely, individual assemblies can outperform co-assemblies in samples where high levels of microdiversity impede contig formation (46-48). Here, we found that co-assembly of all density fractions generated the most medium- and high-quality MAGs, which agrees with two recent SIP metagenomics studies (23, 24). However, we also found that merging binning results from individual fraction assemblies and larger co-assemblies via MAG de-replication provided more medium- and high-quality MAGs than did co-assembly alone. We posit that this approach reaps the benefits of both strategies: it provides higher read recruitment for assembling rare genomes in co-assemblies and also leverages lower microdiversity in individual fraction assemblies. Optimal assembly strategies may differ for other environmental samples, and these strategies must be re-evaluated as sequencing and assembly methods evolve, but our results suggest that SIP metagenomic studies can benefit from employing multiple assembly approaches to maximize genome recovery.

Processing DNA-SIP samples is laborious, but semi-automated protocols simplify lab work and enable high-throughput SIP metagenomic studies (24). Indeed, increasing the number of biological replicates, and sequencing more density fractions per replicate, can improve the detection of labeled taxa (41). However, the opportunities for accidental mistakes, such as cross-contamination, sample mixups, or clerical errors, also increase when processing dozens of samples and hundreds of density fractions. In addition, slight mishandling of ultracentrifuge tubes can disturb delicate CsCl gradients (7), and potentially alter genome distributions along the gradient. If left undetected, these types of accidents could produce inaccurate weighted BD estimates, adding extra noise to the data analysis and even compromising results. In this study, we found that including pre-centrifugation spike-ins, which had distinct and predictable distribution patterns along the gradient, helped us identify and remove problematic samples before they negatively impacted our analyses. Including internal standards can mitigate potential errors and enhance the quality of large complex SIP studies with many replicates. Moreover, with careful design and additional development, internal standards might someday correct for variability introduced during sample processing (41) instead of simply flagging samples for removal. Internal standards can be easily incorporated into automated SIP metagenomics protocols (24), where they can improve the quality of SIP metagenomic results, and if adopted broadly, potentially serve as consistent fiducial reference points that facilitate inter-comparisons of different SIP studies.

Accurate measurements of genome abundance along the BD gradient are essential for identifying labeled genomes and determining their level of isotopic enrichment (30). However, the compositional nature of metagenomic data, and the variability introduced during sample processing and sequencing, can hamper quantitative estimates of genome abundance (25-28, 49). Internal quantification standards can mitigate process variability and provide absolute abundance estimates of genes, transcripts, and genomes from metagenome and metatranscriptome data (28, 37, 50-53). Based on these findings, we hypothesized that adding internal standards to density fractions (“sequins”) could improve abundance measurements and thereby improve isotope enrichment measurements. Indeed, estimates of AFE in our study were more accurate using absolute abundances derived from sequin normalization compared to AFE estimates using other strategies.

Multiple factors could explain the more accurate estimates of isotopic labeling enabled by internal quantification standards. For one, sequins may have mitigated any variation introduced during library creation and sequencing (28). Additionally, sequins may have corrected for differences in DNA recovery among fractions that would have otherwise gone unnoticed and negatively impacted abundance measurements. That is, after collecting CsCl fractions, each fraction separately undergoes PEG precipitation and desalting before DNA concentrations are measured (24). Absolute abundances calculated using DNA concentrations assume identical DNA recovery efficiencies (22, 23), so any stochastic or systematic variability in the percent of DNA recovered would lead to errors in absolute abundance measurements. Conversely, sequins track and mitigate variability in DNA recovery when they are added to fractions before the desalting steps, as was performed here. Therefore, if DNA recovery efficiency varied among fractions, then we would expect absolute abundances derived from sequins to be more accurate than estimates derived from DNA concentration measurements. Without internal standards, variability introduced during DNA recovery, library construction, and sequencing is unknowingly propagated as noise into downstream SIP analyses. This undetected variability can potentially lead to errors that impact predictions of isotope enrichment.

The various SIP analysis methods examined in this study use different approaches to detect labeled microorganisms, and these differences could impact the sensitivity and specificity of their predictions. The accuracy of different SIP analysis methods has not been assessed with metagenomic data until now, but *in silico* simulations of 16S rRNA-based SIP data revealed that MW-HR-SIP had higher balanced accuracy than the other analysis methods (29). The qSIP model also generated more accurate AFE estimates than the ΔBD method in those simulations. We observed similar patterns by comparing analysis methods using our experimentally-designed SIP microbiome. In addition, we found that the consensus of multiple approaches, i.e., MW-HR-SIP and qSIP, produced higher accuracy results than any single method alone. Future SIP metagenomic studies might increase confidence in identifying isotope-incorporating taxa by employing these two independent strategies, although the higher confidence in true positives might come at the cost of missing labeled genomes with lower coverage. Regardless of the analysis tools used, analyzing more biological replicates is another simple strategy to increase accuracy (41). As SIP analysis methods evolve, reassessing their performance with deeper sequencing, more replicates, and an improved mock microbiome (e.g. more species at different AFE levels) will provide additional insights into their accuracy and limitations.

Altogether, we used a first-of-its-kind mock SIP metagenome to assess the performance of different analysis approaches, identified a set of current best practices, and established an experimentally validated workflow for SIP metagenomics. The ‘wet-lab’ aspects of the workflow include the addition of pre-centrifugation spike-ins for quality control and post-fractionation sequins for genome quantitation along the BD gradient. The ‘dry-lab’ aspects entail absolute genome normalization in each density fraction, and a modified qSIP model tailored to handle genome-resolved metagenomic datasets to calculate AFE. We also explored strategies to more accurately identify isotope incorporators, such as limiting analysis to taxa with coverage and isotope enrichment levels above minimal thresholds and using the consensus of multiple SIP analysis tools to detect labeling using our newly developed *SIPmg* package. These additional strategies hold promise for improving the accuracy of SIP metagenomic results, although the specifics of how and when to apply them will depend on the study design and individual preferences regarding the tradeoffs between specificity and sensitivity. We believe this validated analysis workflow will increase the reliability of SIP metagenomic findings, enable standardization across studies, and facilitate the use of SIP data in modeling microbially-mediated processes.

## MATERIALS AND METHODS

### DNA collection and mock community creation

To create a mock microbiome where the identity of labeled genomes and their level of enrichment were known *a priori*, we first extracted DNA from bacterial isolates grown in ^13^C-labeled glucose. *Escherichia coli* K-12 wildtype cells were grown in M9 minimal salts media (Teknova; M8005). Glucose was added at a final concentration of 20mM and was the sole carbon source in both media. DNA with different levels of ^13^C enrichment was produced by varying the ratio of unlabeled glucose to uniformly-labeled ^13^C_6_-D-glucose (Cambridge Isotope Laboratories; CLM-1396; 99 atom %), e.g. DNA extracted from cultures grown in a ratio of 4:1 of unlabeled:labeled glucose was assumed to have an enrichment of approximately 20 atom %. Cultures grown overnight in LB were transferred into labeled media at 5,000-fold dilution (i.e. 2ul into 10ml labeled media), grown at 37°C, and harvested at mid-log phase. DNA was extracted using the Wizard genomic DNA purification kit (Promega; A1120) and quantified using the QuantIT dsDNA High Sensitivity Assay Kit (ThermoFisher; Q33120).

DNA from a complex microbial community was recovered from an outdoor, man-made pond located at the Joint Genome Institute. Pond water was pre-filtered through a 5 um mesh before collection onto 0.2 um Supor filters (Pall; 47 mm dia.). DNA was extracted from filters using a DNeasy PowerWater kit (Qiagen; 14900-50-NF).

Replicate samples were prepared for ultracentrifugation by combining 900 ng of microbiome DNA with 50 ng DNA from each bacterial isolate. For samples with isotopically labeled DNA, the ratio of unlabeled to labeled DNA for each isolate was adjusted, e.g. 40 ng of unlabeled *E. coli* DNA was combined with 10 ng of 20% enriched *E. coli* DNA. The specific ratios of unlabeled:labeled DNA are described in Table S1.

### Synthetic pre-centrifugation DNA spike-ins

A set of six synthetic DNA fragments were added to mixtures of DNA from isolates and the complex microbiome to track the ultracentrifugation and fraction collection steps. These fragments were approximately 2 kbp in length with GC content of 37-63% (Table S2). To change the distribution of fragments across the density gradient, some fragments were artificially enriched with ^13^C through PCR by adjusting the ratio of unlabeled dNTPs and uniformly-labeled ^13^C dNTPs (Silantes Gmhb; 120106100; >98 atom %) (Table S2). Briefly, DNA was amplified for 30 cycles by adding 0.5ul Phusion High Fidelity DNA Polymerase (NEB; M0530S), 10ul of 5X Phusion HF Buffer, 1ul of 10 mM dNTPs (final conc. labeled/unlabeled mixture), 2.5 ul each 10 μM Forward and Reverse Primer, and 31.5 ul of nuclease-free water. PCR products were purified using AMPure XP beads (Beckman Coulter; 63880) and pooled in equimolar ratios to create a set of pre-centrifugation DNA spike-ins. These pre-centrifugation spike-ins were added at 1% by mass of the DNA mixture, e.g. 10 ng of synthetic fragment pool added to 1 ug of microbial DNA mixture.

### Gradient separation, sequin addition, and fraction purification

Following Nuccio and colleagues (24), samples were centrifuged at 44,000 RPM (190,600 g) for 120 hours at 20°C in a VTi 65.2 Rotor (Beckman Coulter; 362754). For each sample, 24 fractions of 220 uL were collected into a 96-well plate using an Agilent 1260 fraction collector running at flow rate 250 uL/min while using mineral oil as the displacement fluid. Fraction density was determined using a Reichert AR200 refractometer.

Before purifying DNA from CsCl fractions, an additional set of 80 synthetic DNA fragments, or *sequins* (28), were added to each fraction as an internal standard for subsequent quantitative metagenomic analysis. Lyophilized pellets of sequins were obtained from the Garvan Institute of Medical Research (https://www.sequinstandards.com). Pellets were resuspended in TE Buffer (10 mM Tris, 0.1 mM EDTA, pH 8.0), and the concentration was measured with QuantIT dsDNA High Sensitivity Assay Kit (ThermoFisher; Q33120). Of the 24 BD fractions collected for each sample, we selected 16 to move forward with library creation and sequencing based on the range of BD they spanned. These 16 fractions were amended with sequins. To compensate for expected differences in the amount of DNA recovered from different densities, the middle 8 fractions received 300 pg of sequins while the 4 fractions on either tail received 100 pg of sequins.

After sequin addition, DNA was recovered by adding a 250 ul solution of 36% 6000 PEG and 1.6M NaCl to each fraction and incubating overnight in 4°C. Plates were centrifuged at 3,214 x*g* for 1.5 hours at 20°C to pellet DNA. Pellets were washed with 300 ul of 70% chilled ethanol, centrifuged at 3,214 x*g* for 45 minutes at 20°C, and resuspended in 30 ul of TE Buffer (10 mM Tris, 0.1 mM EDTA, pH 8.0). Purified DNA was quantified using Quant-IT dsDNA High Sensitivity Assay Kit (ThermoFisher; Q33120).

Sequins were added to each fraction before PEG precipitation and DNA quantification steps; therefore the amount added was based on the expected sample DNA concentrations. Tailoring sequin additions to actual sample DNA concentrations, as opposed to estimates, is preferable to ensure optimal coverage in sequencing data. After completing analysis of the mock microbiome, we sought to improve sequin additions by measuring DNA levels before PEG precipitation when DNA was still in concentrated CsCl. Additional details are provided in the Supplementary Materials.

### Library creation and sequencing

Sequencing libraries were generated from the 16 middle fractions of each sample using Nextera XT v2 chemistry (Illumina) with 12 PCR cycles. Concentrations and size distributions of each library were determined on a Fragment Analyzer (Agilent). Libraries were pooled at equal molar concentrations within the range of 400-800 bp, and the pool was size selected to 400-800 bp using a Pippin Prep 1.5% agarose, dye-free, internal marker gel cassette (Sage Science). For each library, 2×150 bp paired-end sequencing was performed on the Illumina Novaseq platform using S4 flowcells (Table S7).

### Metagenome assembly and binning

Raw reads were filtered and trimmed using RQCFilter2 software according to the standard JGI procedures (https://jgi.doe.gov/data-and-tools/software-tools/bbtools/bb-tools-user-guide/data-preprocessing/). Then, one of the four strategies was used to perform contigs assemblies: a) an assembly of unfractionated SIP sample with metaSPAdes(v3.15.2) (54); b) a single fraction assembly with metaSPAdes (371 assemblies); c) a single sample co-assembly with metaSPAdes (co-assembly of all fractions sequenced for a single SIP replicate sample, 24 assemblies); d) an experiment-wise co-assembly with MetaHipMer(v.2.0.1.2) (assembly of all fractions across all replicates) (32). Assembly and genome mapping parameters are reported in the Supplementary Methods. We generated 397 assemblies in total. Quality assessment metrics for each assembly were calculated using QUAST(v5.0.2) (MetaQUAST mode)(Data Set S3) (55). Each assembly was then independently binned with MetaBAT(v2.12.1) (56). For each generated MAG, we used GTDB-Tk(v2.0.0) (GTDB R95) (57) to assign a taxonomic classification. To assess the quality of MAGs we used CheckM(v1.1.3) (58) and QUAST(v5.0.2) (59). The MetaHipMer combined assembly was annotated using the JGI metagenome annotation workflow (56) and is available through IMG/M (60) under taxon identifier 3300045762.

### MAG deduplication and mean scaffold coverage calculations

Medium- and high-quality MAGs recovered from all assembly strategies were deduplicated to remove redundant versions of each draft genome (34). The genome-wide ANI (gANI) and the alignment fraction (AF) were calculated for each possible MAG pairwise comparison (35). Next, the lowest pairwise values of gANI and AF were utilized for each MAG comparison, followed by clustering using single-linkage to group MAGs based on species-level delineations (e.g., gANI >= 96.5 and AF >= 30) as defined by Varghese and colleagues (35). MAGs that did not cluster with other MAGs were considered singletons. Following clustering, we used completeness, contamination, and total length values to select a single representative MAG for each cluster. Sequences of all spike-ins and sequins were concatenated with the final set of MAG contigs, and this contig set was then used as a reference for read mapping across all density fractions (see Supplementary Methods). The average contig coverage of MAGs, spike-ins, and sequins in each fraction was calculated and used in the downstream analysis.

### Quality control of SIP data using pre-centrifugation spike-ins

Before performing SIP analysis, we first removed mishandled samples from our dataset. For this purpose, we identified the peak of absolute concentration distributions across the density gradient for each labeled pre-centrifugation spike-in. If the spike-in distribution patterns did not match the expected order along the density based on the theoretical estimated density of the spike-in (given its GC content and C^13^/C^12^ ratio), then the sample was considered potentially problematic and removed from the analysis.

### Estimating the absolute abundance of MAGs across density fractions

To determine the extent of isotope incorporation into genomes, it is first necessary to measure genome abundance across the density gradient. We explored several ways to measure genome abundance in the SIP dataset, which are implemented as part of the *SIPmg* R package (see Code Availability).

First, we obtained absolute concentrations of genomes across the density gradient using the approach proposed by Hardwick and colleagues (28), in which sequins were used as internal reference standards to scale coverages into absolute concentrations. Briefly, the average MAG coverage within a given fraction (metagenome) was scaled into units of molarity using regression analysis based on known molarity of 80 sequins and their average coverages. Molar concentrations of the sequins in the added standard mixture were obtained from the manufacturer (Garvan Institute of Medical Research). For regression analyses, we first tested both ordinary least squares regression and robust linear regression. When using ordinary least squares regression, we also tested Cook’s distance filtering to remove outliers at a threshold of Cook’s distance < n/4 (n is the number of datapoints in the regression analysis). A coefficient of variation threshold of 250 was employed as a quality control step in this scaling process. Due to the lower number of false positives in the approach with ordinary least squares regression combined with Cook’s distance filtering, we continued with this approach for all analyses, but also report the findings from using the robust linear regression analysis in the Table S3. A detailed workflow for sequin normalization is provided in the vignette for the *SIPmg* R package (https://github.com/ZielsLab/SIPmg).

In addition to sequin based normalization, we also explored genome abundance estimation using: (1) unscaled coverage; (2) relative coverage; (3) absolute abundance as per the approach of Greenlon and colleagues (23) and as the per approach of Starr and colleagues (22). Unscaled coverages represented raw average MAG coverage values that were directly used in the estimation of mean weighted BDs and AFE. Relative coverage was estimated as: (coverage of a MAG within a fraction)/(sum of coverages of all MAGs within a fraction).

### Estimating of atom fraction excess of MAGs

The qSIP model (eq. 1) or ΔBD model (eq. 6) can be used to estimate the AFE of genomes. Briefly, the AFE of organism *i*, can be quantified using the qSIP approach (30):

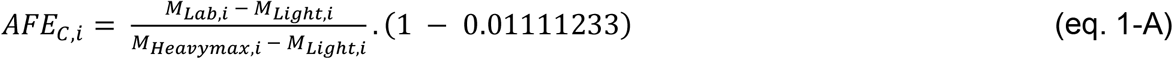

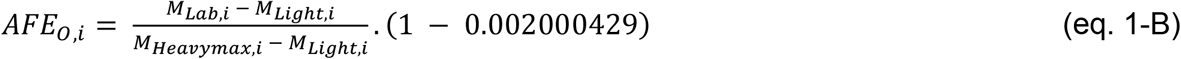

where: *A*_*C,i*_ and *A*_*O,i*_ are the estimated AFE with oxygen and carbon as the isotopic substrate, respectively. *M*_*Light*_ is the molecular weight of a MAG (g/mole) in the control condition (eq. 2), *M*_*Lab*_ is the molecular weight of a MAG (g/mole) in the treatment condition (eq. 3), and *M*_*Heavymax*_ is the theoretical maximum molecular weight of a MAG (g/mole) due to the maximum labeling by the heavy isotope (eq. 4) in the treatment condition:

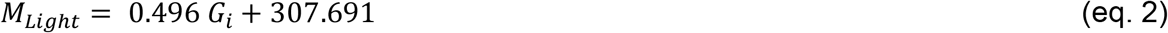

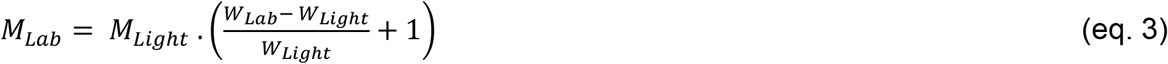

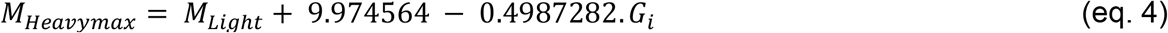

where: *G*_*i*_ is the GC content of the MAG (ranging from 0 to 1). Here, we modified the qSIP model to use the GC content values of MAGs provided from output of CheckM (58), rather than inferring it using an empirical regression (30). *W*_*Light*_ and *W*_*Lab*_ are the mean weighted buoyant densities across control and treatment conditions respectively.

The weighted average buoyant density (W_ij_) is then estimated as:

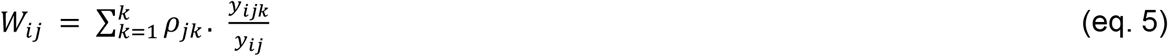

where: *ρ*_*jk*_ is the buoyant density of fraction *k* in replicate *j, y*_*ijk*_ is the absolute concentration of taxon *i* in fraction *k* of replicate *j*, and *y*_*ij*_ is the sum total of absolute concentration of taxon *i* in replicate *j*. Here, genome abundances were determined using either (1) sequin normalization; (2) relative abundance per coverage and/or reads mapped multiplied by total DNA concentrations; and (3) relative coverage.

The estimation of AFE based on the ΔBD model can be represented as:

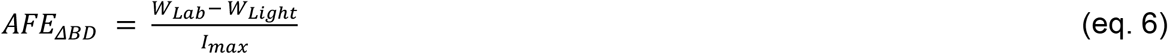

where: *I*_*max*_ is the maximum linear shift in DNA BD (upon 100% labeling), as discussed by Birnie and Rickwood (61). The weighted mean BDs were the same as estimated from eq. 5. This is a variant of ΔBD from the Pepe-Ranney and colleagues study (8), in which OTU read counts were interpolated at specific points of the replicate BD gradients to estimate weighted mean BDs. The above models for determining AFE were incorporated into the *SIPmg* R package for application with SIP metagenomics datasets.

### Identifying isotope incorporators using HR-SIP and MW-HR-SIP

To run the HR-SIP and MW-HR-SIP methods, we used the MAG abundances obtained from the sequin normalization approach. Differential abundances based on absolute abundance for MAGs in the heavy fractions in the treatment conditions were compared to control conditions using HR-SIP and MW-HR-SIP using the HTSSIP R package (29). For HR-SIP, a heavy BD window was set from 1.71 g/mL (as the theoretical peak of *E. coli* would be at 1.709 g/mL based on a GC content of 0.504) to the maximum buoyant density in every treatment condition. For MW-HR-SIP, the overlapping heavy buoyant density windows chosen were 1.71 - 1.74 g/mL, 1.72 - 1.75 g/mL, and 1.73 - 1.76 g/mL. In all cases, sparsity thresholds between 0% and 30% at 5% intervals were chosen (e.g., sparsity threshold of 25% maintains that MAGs must be present in >25% of fractions in the testing windows). The sparsity threshold with the greatest number of rejected hypotheses were selected for final inference of incorporator identity. The Benjamini-Hochberg method was used to adjust for multiple testing with a threshold of *p*-value of 0.05 to identify incorporators.

### Subsampling of *E. coli* reads

Reads that mapped to *E. coli* MAG were extracted from .bam files and subsampled using samtools (v1.7) (htslib 1.7) at 10, 1, 0.1, 0.01, and 0.001 percentages. New *E. coli* MAG coverages for each fraction were then calculated (Table S5) and used in SIP analysis to establish limitations that low coverage input may have on the efficiency of bacterial incorporator identification.

## Data availability

Raw metagenome sequencing reads have been deposited under BioProject Accession PRJNA878529. The MetaHipMer combined assembly and annotated data is available through IMG/M under taxon identifier 3300045762. Single-fraction and combined per-sample assemblies, along with all MAGs and input files for qSIP analysis are available via https://portal.nersc.gov/dna/microbial/prokpubs/DVyshenska2022/. A full list of available data and associated NCBI accession numbers are available in Data Set S3.

## Code availability

The code for the *SIPmg* R package is available for download, along with a vignette describing all functions, at: https://github.com/ZielsLab/SIPmg. The *SIPmg* package includes functions to calculate global scaling factors for genomes based on regression of sequin coverage versus concentration using either ordinary least squares linear regression or robust linear regression. The package can thereafter estimate AFE using either qSIP model or ΔBD method. The package also outputs both FCR adjusted and Bonferroni adjusted bootstrapped AFE confidence intervals for MAGs. The package can also perform HR-SIP and MW-HR-SIP which were built using the HTS-SIP R package.

## ACKNOWLEDGMENTS

The work (proposal: 10.46936/10.25585/60001104) conducted by the U.S. Department of Energy Joint Genome Institute (https://ror.org/04xm1d337), a DOE Office of Science User Facility, is supported by the Office of Science of the U.S. Department of Energy under contract no. DE-AC02-05CH11231. P.S. was supported by a Discovery Grant to R.Z. (RGPIN-2018-04585) from the Natural Sciences and Engineering Research Council of Canada (NSERC). E.E.N., S.J.B. and J.P.R. were supported by a JGI Emerging Technologies Opportunities Program award (DOI: 10.46936/10.25585/60008401) and DOE award SWC1632. Work at Lawrence Livermore National Laboratory was conducted under the auspices of the U.S. DOE under Contract DE-AC52-07NA27344. We thank Tim Mercer at the Garvan Institute of Medical Research for providing sequins.

## AUTHOR CONTRIBUTIONS

E.F., R.M., R.Z., E.E.N, S.J.B, J.P.R, and D.V. conceived the study. D.V. was in charge of overall direction and planning. K.S., A.T., E.E.N., and W.B.K carried out laboratory work. R.Z, P.S., and D.V. developed the analytical strategies for utilizing internal standards for quantitative SIP metagenomics, and D.V and P.S performed the computational analysis. P.S. developed *SIPmg* R-package. A.C. and R.R. performed MetaHipMer co-assembly. N.V. performed metagenome binning and ANI-AF computation. S.R. contributed to the interpretation of the results. M.K. supported metagenome binning coverage analysis. D.V. and P.S. wrote the manuscript. E.F, R.M., and R.Z. supervised the study. All authors provided critical feedback and helped shape the research, analysis, and manuscript.

## SUPPLEMENTAL MATERIAL FILE LIST

**Table S1**. *E. coli* AFE (%) in each treatment condition.

**Table S2**. Characteristics of pre-centrifugation spike-ins. To produce distinct distribution patterns along the density gradient, some spike-ins were artificially enriched with ^13^C through PCR by adjusting the ratio of unlabeled dNTPs and uniformly-labeled ^13^C dNTPs. Theoretical AFE values are reported based on the ratio of labeled dNTPs, but actual AFE values were not experimentally confirmed.

**Table S3**. Comparison of various abundance estimation strategies. All results were derived from the qSIP analysis method. Sensitivity and specificity were averaged across the seven treatment conditions.

**Table S4**. Comparison of methods to identify isotopically labeled genomes. Evaluations were based on absolute genome abundances obtained by normalizing coverage to internal sequin standards using the sequin approach. Specificity and sensitivity were averaged across the seven treatment conditions.

**Table S5**. Average total coverage across all fractions for *E. coli* in different treatment conditions after subsampling from 100% to 0.001% of the original *E. coli* sequence reads.

**Table S6**. Comparison of MW-HR-SIP and qSIP methods for detecting isotopic labeling of *E. coli* at different levels of total genome coverage across the density gradient. ‘True’ indicates *E. coli* was correctly identified to be isotopically labeled (true positive), and ‘false’ indicates *E. coli* was incorrectly identified as unlabeled (false negative). NA corresponds to the failure of the MW-HR-SIP algorithm with that dataset.

**Table S7**. The impact of genome coverage levels on detecting isotope incorporation using the qSIP model.

**Table S8**. Comparison of MAGs retained and the number of false positives detected using the qSIP method after applying different minimum genome coverage thresholds. MAGs were retained if their average total coverage in the unlabeled controls exceeded the coverage threshold. *E. coli* was the only true positive and had a coverage of 1029X, thus no false negatives were detected using the coverage thresholds below.

**Table S9**. Comparison of specificity, sensitivity, and balanced accuracy of the qSIP method after applying minimum AFE thresholds. To be identified as isotopically labeled, the lower 95% CI interval of a genome’s estimated AFE must be greater than the minimum AFE threshold.

**Table S10**. Comparison of false positives MAGs identified by the MW-HR-SIP, qSIP, and ΔBD methods. Names of the false positive MAGs are listed in each column.

**Table S11**. Comparison of *E. coli* AFE confidence intervals estimated using qSIP alone, qSIP after first applying MW-HR-SIP, and qSIP after first applying ΔBD method to identify a subset of putatively labeled MAGs. Condition B (“20pct_20ng”) was removed as it *E coli* was never identified as an isotope incorporator in this condition.

**Figure S1**. Average completeness and average purity of MAGs grouped by assembly type (I - intact metagenome assembly with MetaSPAdes, F - separate fractions assembled with metaSPAdes, S - all fractions within each replicate co-assembled with metaSPAdes, M - combined assembly of all fractions using MetaHipMer(v.2.0.1.2))

**Figure S2**. Average coverage across all fractions for each medium and high-quality MAG. Color-coding identifies MAGs found in multiple assembly types (Shared) or uniquely generated in one of the three different assembly types (F - separate fractions assembled with metaSPAdes, S - all fractions within each replicate co-assembled with metaSPAdes, M - combined assembly of all fractions using MetaHipMer). Assemblies of unfractionated DNA (Intact) with MetaSPAdes did not generate unique MAGs.

**Figure S3**. Detecting anomalous samples using pre-centrifugation spike-ins. A) SIP sample displaying the expected spike-in distribution pattern based on relativized absolute coverage along the density gradient. B) An anomalous sample whose spike-in patterns do not match expectations, indicating possible problems in gradient collection and library creation.

**Figure S4**. Linear regression parameters and Spearman correlations between estimated and expected AFEs obtained using the modified qSIP model from (a) raw coverage, (b) relative coverage, (c) multiplying relative abundance with DNA concentration following Greenlon and colleagues (23), (d) multiplying relative coverage with DNA concentration following Starr and colleagues (22), (e) Sequin approach with ordinary least squares regression without Cook’s distance filtering (f) Sequin approach with ordinary least squares regression with Cook’s distance filtering (g) Sequin approach with robust linear regression, and (h) Relativizing abundances per fraction (MAG abundance/sum of MAG abundances in each fraction) from sequin approach with robust linear regression. *p*_*reg*_ and *p*_*cor*_ correspond to the *p-*values for the linear regression and Spearman correlation. The intercepts determined by linear regression were not significantly different from zero (*p-value* > 0.05) in any method for estimating abundance.

**Figure S5**. Linear regression parameters and Spearman correlations between estimated and expected AFEs obtained using the ΔBD method from (a) raw coverage, (b) relative coverage, (c) multiplying relative abundance with DNA concentration following Greenlon and colleagues (23), (d) multiplying relative coverage with DNA concentration following Starr and colleagues (22), (e) Sequin approach with ordinary least squares regression without Cook’s distance filtering (f) Sequin approach with ordinary least squares regression with Cook’s distance filtering (g) Sequin approach with robust linear regression, and (h) Relativizing abundances per fraction (MAG abundance/sum of MAG abundances in each fraction) from sequin approach with robust linear regression. *p*_*reg*_ and *p*_*cor*_ correspond to the *p-*values for the linear regression and Spearman correlation. The intercepts determined by linear regression were not significantly different from zero (*p-value* > 0.05) in any method for estimating abundance.

**Figure S6**. Linear regression parameters and Spearman correlations between estimated and expected AFEs obtained using the qSIP method for subsampled data at mean cumulative coverages of (a) 0.01X, (b) 0.1X, (c) 1X, (d) 10X, (e) 100X, and (f) 1000X. *p*_*reg*_ and *p*_*cor*_ correspond to the *p-*values for the linear regression and Spearman correlation. The intercepts determined by linear regression were not significantly different from zero (*p-value* > 0.05) at any level of subsampling.

**Figure S7**. Mean total coverage of MAGs across biological replicates in the unlabeled controls. False positive MAGs are indicated by blue bars (also indicated by arrows). The mean coverage threshold where false positives would be removed (17X) is indicated by a dashed horizontal line. A total of 68 MAGs had mean total coverages greater than this threshold. MAGs lower than this threshold are separated by a dashed vertical line.

**Figure S8**. Mean specificity of delta BD, modified qSIP, and MW-HR-SIP methods to infer incorporators. The error bars indicate standard deviation of specificity across the seven treatment conditions. The annotations on the bars indicate the number of false positives out of 146 MAGs.

**Figure S9**. Impact of SIP CsCl gradient solution on measurements of DNA concentrations made with the Quant-IT DNA High Sensitiviy Assay Kit. The error bars indicate standard deviation (n=5). The dashed line indicates a linear regression (R^2^=0.9875; F-test *p-value* = 6.32 × 10^−8^).

**Data Set S1**. Internal calibration standards utilized in experimental design. A set of six synthetic DNA fragments (pre) were added to mixtures of DNA from isolates and the complex microbiome to track the ultracentrifugation and fraction collection steps. An additional set of 80 synthetic DNA fragments (post), or sequins, were added to each fraction as an internal standard for subsequent quantitative metagenomic analysis.

**Data Set S2**. Metagenome-assembled genomes (MAGs) generated across assembly approaches and associated quality metrics. A total of 2,022 MAGs were generated across all assemblies, of which 248 were high-quality, 447 were medium-quality, and 1,327 were low-quality as defined by the MIMAG reporting standards. Bin identifiers and assembly identifiers are provided, along with CheckM metrics for estimates of completeness and contamination. Cluster representatives are denoted based on single-linkage clustering from average nucleotide identity values of ≥ 96.5 and alignment fractions of ≥ 30%.

**Data Set S3**. Metagenome assembly types, metrics, and associated accessions for GOLD and NCBI.

